# Plant growers’ environmental consciousness may not be enough to mitigate pollinator declines: a questionnaire-based case study in Hungary

**DOI:** 10.1101/2022.04.07.487523

**Authors:** Zsófia Varga-Szilay, Gábor Pozsgai

## Abstract

**BACKGROUND:** Pesticides are one of the most important anthropogenic-related stressors. In times of global pollinator decline, the role of integrated farming and that of urban gardens in supporting wild pollinators is becoming increasingly important. We circulated an online questionnaire to survey the plant protection practices among Hungarian farmers and garden owners with a particular emphasis on pollinator protection.

**RESULTS:** We found that plant growers heavily rely on pesticide use, and pesticides are widely used in otherwise pollinator-friendly gardens. Whether pesticide use practices were driven by expert opinion and the respondents’ gender were the best predictors of pesticide use. Although most respondents supported pollinators, pesticides are also widely used among home garden owners, which can pose a non-evident ecological trap for pollinator populations in the gardens.

**CONCLUSION:** Special attention should be paid to implementing measures to reduce pesticide, use not only in farmlands but also in home gardens. Environmental education and financial support through agroecological schemes could efficiently promote the transition. However, whereas farmers can be encouraged to reduce pesticide use mostly by expert advice, garden owners are likely to rely on more conventional information channels. The attitude of Hungarian plant growers can provide an insight into pesticide use practices of Central and Eastern European countries, but similar surveys are needed across Europe for a complete understanding of broad-scale processes. This work lays the foundations for similar studies which can inform and facilitate the transformation processes to pesticide-free farming and gardening.

## 1 INTRODUCTION

In the Anthropocene, biodiversity declines at an alarming pace. One of the important groups, the insects, is among the most impacted (e.g. Hallmann et al. ^1^; Zattara and Aizen ^2^). Insect declines pose a major threat to a variety of ecosystem functions and the delivery of the derived ecosystem services, all of which are vital to humans ^3,4^. One important ecosystem service, mostly provided by insects, is pollination ^5^. Insect pollinators suffer from habitat fragmentation, reduction of flower resources, lack of nesting space, as well as from exposure to pesticides from agricultural activities ^6–8^. Despite their long-known negative effects on human and environmental health ^9,10^, pesticides are widely used both in industrial-scale farming and urban green areas and their application has even increased with agricultural intensification in recent decades ^11^. Indeed, the spillover of chemical insecticide residues from farmland can negatively affect wild insect pollinators in adjacent natural and semi-natural areas ^12,13^ causing direct mortality, behavioural abnormalities, and reduced reproduction rates ^14^. Furthermore, the concomitant use of agrochemicals (pesticides and fertilisers) can cause an even more detrimental ‘cocktail effect’ to insect pollinators ^15–17^. In fact, a combination of over sixteen different agrochemicals was detected in flying insects in nature conservation areas adjacent to agricultural lands across Germany ^18^ and the USA ^19^. Thus, agrochemicals are suggested to play a major role in driving global insect declines ^20,21^, particularly on farmlands ^22^.

To address biodiversity loss on farmlands, particularly that of pollinators, the European Commission created a farm strategy to cut the use of chemical pesticides in European countries ^23^ and the reduction or complete elimination of pesticide use has been advised by the scientific community (e.g. Goulson and Nicholls ^7^). A number of modern synthetic pesticides have been banned (e.g. in the European Union all neonicotinoids except acetamiprid) after they have been proven to harm non-target insects (like bees) in addition to the pest species targetted. In fact, the transition to alternative agricultural practices is possible without yield losses ^24^ whilst pest damage can be reduced and farm profitability maintained after lowering, but not completely abandoning, pesticide use ^25,26^. Despite the increasing number of organic farms ^27,28^ in the European Union, which may be the first step toward a pesticide-free, and thus a biodiversity-friendly, farming, the conversion process can take years because the current conventional plant protection strategies employed on non-organic farms still require synthetic pesticide input. Nevertheless, evidence suggests that these integrated efforts may be a first step toward maintaining healthy ecosystems. For example, management that promotes ecosystem services (such as biological control or pollination) can support high insect diversity in areas of agricultural mosaics ^29^. Moreover, even in conventionally managed farms, increasing the proportion of semi-natural habitats, such as hedges or field-edge flower strips, can dramatically increase the diversity of insects that are beneficial to agriculture, including that of pollinators ^30,31^. However, although increasing natural habitat areas or employing other integrated pest management approaches can lead to increased pollinator and other insect diversity, unless these ecosystem-based approaches are combined with pollinator-friendly management, their positive effect will be reduced or completely eliminated by the use of synthetic pesticides ^32–35^. Since socio-economic factors can dictate how rapidly the transition to pesticide-free farming unfolds, knowing farmers’ approaches to these novel strategies is essential for future planning.

Whilst it may be difficult to achieve pesticide-free pest control within high-intensity farming (especially in monocultures), it may be a more feasible approach in small-scale farms and urban areas. Small-scale sustainable farming systems and well-planned urban green areas, such as biodiversity-friendly parks and allotments (community gardens), can mitigate pollinator declines ^36,37^. In fact, in a landscape mosaic with a high proportion of urban areas, organically managed parcels of land can maintain high biodiversity and serve as a source of native pollinators within a landscape where most land is not managed with the maintenance of biodiversity as a key goal ^38,39^.

Moreover, an increasing number of scientific papers support the premise that urban and suburban gardens function as refuges and local hotspots for biodiversity ^40,41^, and support diverse communities of insect pollinators, even in highly urbanised areas ^42^. These gardens can be near-natural and support viable metapopulations of rare species ^43^. However, the true conservation potential of human-altered areas for pollinators depends on the available floral resources, nesting and hiding spaces, and on the proportion of near-natural areas that can be found in the urban landscape. These factors also determine the abundance and diversity of pollinator communities ^37,44^.

Urban gardens may not be always beneficial for insect pollinators though. First, there is a wide selection of pesticides in shops and supermarkets that are targeted at domestic users and which may be applied in otherwise pollinator-friendly gardens. Second, synthetic pesticides can also get into gardens unintentionally when ornamental plants sold as ‘bee-friendly’ in horticultures are treated with various fungicides and insecticides ^45^. As a consequence, insects lured to supposedly pollinator-friendly gardens can be exposed to a number of synthetic pesticides (especially neonicotinoids) and their residues and this exposure, in turn, can lead to lethal and sublethal effects ^19,46^. The process of banning synthetic pesticides for non-agricultural uses has already begun in some European countries (such as France ^47–49^ but others, including Hungary, are lagging behind. Yet, we have no information on what proportion of private gardens are treated with chemical plant protection products.

There is a large knowledge gap in our understanding of how efficiently farmlands and urban and suburban gardens mitigate insect biodiversity loss at a country scale and how farmers and garden owners approach the transition away from the use of pesticides. Gaining insight into their management habits, motivations and willingness to change is vital for developing further action.

Thus, we conducted a survey to measure plant growers’ dependence on pesticides (highlighting acetamiprid-containing insecticides), particularly to investigate the pesticide application practices and the attitude towards protecting wild pollinators of those who own less than one hectare of land (henceforth home gardens or gardens). We focussed our work on Hungary, a typical, Central-Eastern European country with mainly conventional agriculture in which chemical and more hazardous pesticide use trends are likely to reflect those of general Europeans ^50^.

We aimed to investigate 1) what factors best predict pesticide use in agricultural areas and to what extent plant growers think their application is necessary, 2) to what extent plant growers think the use of insecticides (as a subset of pesticides) is necessary and what are the main considerations determining their selection, 3) how dependent plant growers are on the single currently allowed neonicotinoid (acetamiprid), and 4) if acetamiprid is used, what other pesticides are used simultaneously. Our additional aims were to specifically investigate the home garden owners’ approach to pesticide use and pollinator support. We were interested in 1) how necessary garden owners think it is to use pesticides, 2) what they think about the threats to wild pollinators and how this affects their management practices and 3) what factors predict whether or not gardeners provide support for wild pollinators and what the most common such forms of support are.

We hypothesised that Hungarian plant growers are highly dependent on pesticide input and home garden owners have little awareness of linked environmental issues. Nevertheless, we predicted that home garden owners who predominantly produced for their own needs were more aware of the environmental hazards of pesticides than large-scale farmers and we also hypothesised that the pesticide use among those who supported pollinators was less frequent.

## 2 MATERIAL AND METHODS

### 2.1 Questionnaire Design

We circulated an online questionnaire that consisted of 61 closed-ended questions, all of which were mandatory to respond to. The questionnaire had eight sections to collect information about 1) sociodemographic factors, 2) type of farming, 3) use of plant protection products and 4-6) insecticides and their means of application, 7) protection of wild pollinators, and the questionnaire included one question 8) about how the questionnaire reached respondents (**Supporting Information S1**). All responses were recorded anonymously, however, respondents could provide their email addresses at the end. The questionnaire was designed in Google Forms and circulated in Hungarian language. The questionnaire was shared on social media platforms (such as Facebook groups and Facebook pages, and agricultural websites) and on farming and entomological mailing lists. The form was available from 26 April to 20 August in 2021.

Respondents who do not farm in Hungary were excluded from the analysis and data from Pest county were merged with those from Budapest. The number of respondents was standardised for 100,000 inhabitants in Hungary to improve representativeness.

In this study, we include both chemical and non-chemical pesticides in the group of ‘pesticides’ and ‘plant protection products’. We also use the word ‘insecticide’ inclusively for synthetic insecticides and insecticides that can be used in organic farming.

### 2.2 Data processing and statistical analysis

The original categorical replies were on a few occasions re-categorised for analytical purposes. Education categories were merged into ‘elementary’, ‘middle’, ‘high’, and ‘postgraduate’ levels. The most important sociodemographic parameters and the categories used are listed in **Table 1**, and all other parameters can be found in **Supporting Information S1**. Although they were separated in the original questionnaire, the two kinds of agricultural experts, ‘plant doctors’ and ‘plant protection experts’ were later merged into a combined ‘expert’ category. When the additional plant protection products which were used with acetamiprid were named, they were assigned into nine categories or the combination of those, as ‘adhesion promoter’, ‘insecticide(s)’, ‘insecticide(s) and acaricide(s)’, ‘insecticide(s) and fungicide(s)’, ‘insecticide(s), fungicide(s) and fertiliser’, ‘fertiliser’, ‘fungicide(s)’, ‘fungicide(s) and acaricide(s)’, ‘fungicide(s) and fertiliser’. In the question about how respondents support pollinators the textual responses for food and habitat provision-related answers may overlap although when categorising these, we choose the one which was most strongly emphasised by the respondent. In the same question, we did not create a separate category for ‘outreach’, because it only occurred in a single response. When textual responses were given to the types of support which could not have been categorised as direct action (e.g. ‘I do not harm them’), they were interpreted as ‘no support’.

**Table 1.**
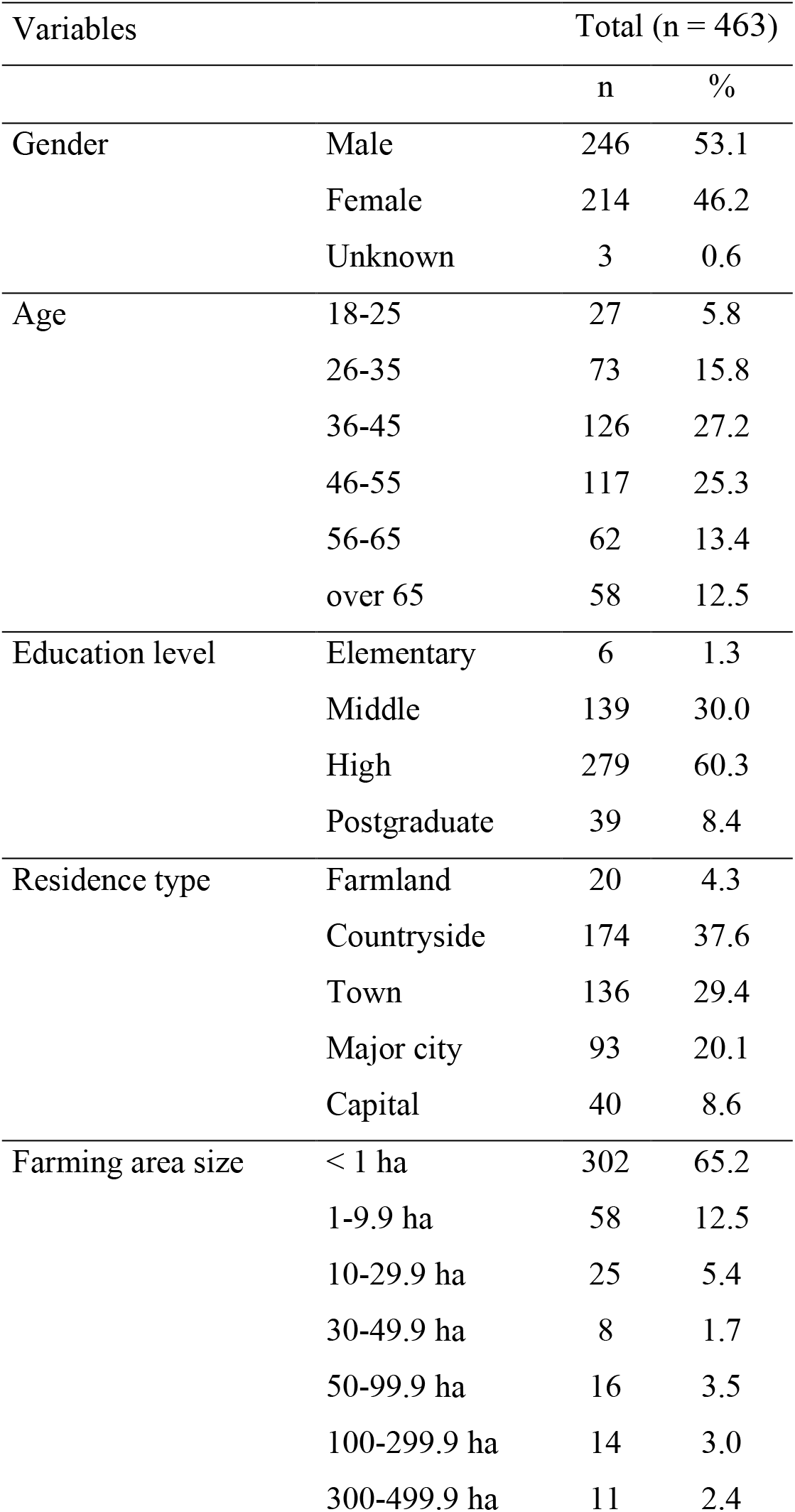
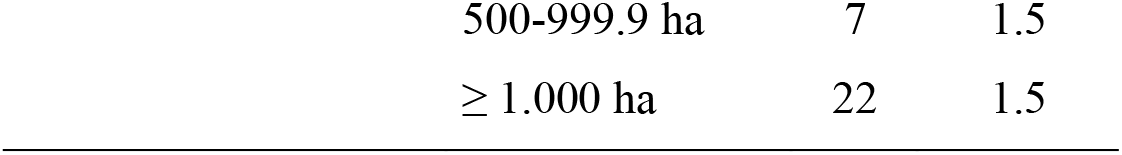
Sociodemographic characteristics of the study population (n = 463)

When the approach solely of garden owners (as a subset of all plant growers) to pesticide use and their attitude to wild pollinators were investigated, only landowners with less than 1-hectare land were included in the analysis.

We used Spearman correlation tests to examine if sociodemographic factors and farming habits correlate with pesticide use. P-values were corrected according to Holm’s method. For calculating the correlation matrix the ‘*psych*’ (version 2.1.9) ^51^, and for visualising them the ‘*corrplot*’ (version 0.92) ^52^ R packages were used. We used the chi-square test to compare plant protection habits of home garden owners and large-scale farmers. We used machine learning techniques with Gradient Boosting Machine (GBM) for generating our models to investigate 1) which socioeconomic factors determine whether or not pesticides were used in farmlands, and 2) which socioeconomic factors determine whether or not pollinators were supported in home gardens. The model fit was evaluated using the Area Under the Curve (AUC) score and by examining the accuracy of the best fitting model. We used the ‘*gbm*’ (version 2.1.8) ^53^, ‘*caret*’ (version 6.0.90) ^54^, and ‘*yardstick’* (version 0.0.9) ^55^ R packages for modelling. Likert Scales figures were plotted using the ‘*likert*’ (version 1.3.5) ^56^ and the map was created using the ‘*sf*’ (version 1.0.3) ^57^ R packages. All analyses were done and figures were created using the R 4.1.1 statistical software ^58^.

## 3 RESULTS

Of the 463 people who completed the questionnaire, 246 were male, 214 were female, and three did not state their gender. The willingness to respond was slightly unbalanced, as more responses were received from the western than from the eastern counties (**Figure 1**). Pest country was the region that yielded the largest proportion of responses (22.0%).

**Figure 1.**
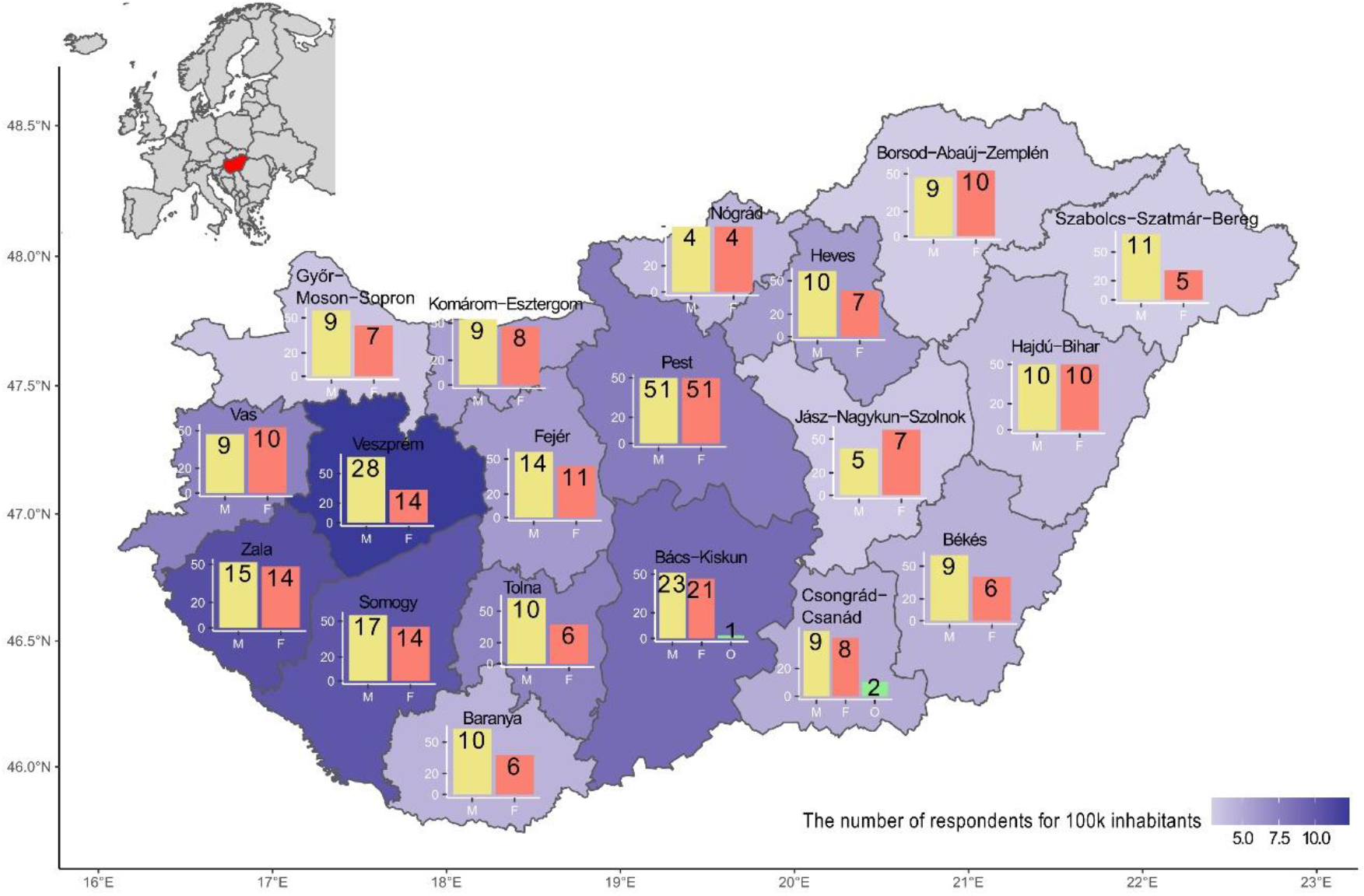
Distribution of respondents by gender (“M” = male, “F” = female, “O” = unknown). Y-axis shows the percentage of the genders and the numbers on the barplots indicate the exact number of respondents in the 19 counties of Hungary. County names are indicated in bold. The colour depth in the map indicates the number of respondents per 100,000 inhabitants. Note that the sum of the numbers indicated on the barplots is greater than the number of respondents because respondents who farmed in more than one county were counted multiple times.

Among the respondents, the two middle-age categories (36–45 and 46–55 years old) were the most frequent, and 60.3% of all respondents (n = 279) fall into the high-level (but not Ph.D.) education group. Of all plant growers, 302 (65.2%) had less than one hectare of farming area (**Table 1**). The most commonly grown crops were vegetables and fruits, followed by grapes and root/tuberous plants (**Supporting Information S2**). Of the respondents, 181 (39.1%) used a pest forecasting system and 370 growers (79.9%) supported natural enemies of pests (**Supporting Information S3**).

### 3.1 Plant protection habits of all plant growers

The majority of plant growers in an area of less than one hectare were individual farmers who produce exclusively for their own consumption (n = 251), while the majority of farmers in an area larger than one hectare either produce for sale privately or as part of a farmers’ association (n = 107). Of these smallholders, over 95% used pesticides (**Table 2**). However, of all respondents, 311 (67.2%) used pesticides, and 212 of them (68.2%) used them together with some additives. Among the pesticide users, 244 (78.5%) usually did not spray during daytime in a flowering culture (**Supporting Information 4**). Of those plant growers who used pesticides, 243 (79.0 %) felt these products were necessary for farming, with 150 (48.2%) of users considering pesticides as being crucial, and 93 (29.9%) of them regarding them as important.

**Table 2.** Distribution of farming types among the study population (n = 463)

| Area size | Type of farming | Uses pesticide |  | Pesticide-free |  |
| --- | --- | --- | --- | --- | --- |
|  |  | n | % | n | % |
| <1ha | Individual: production for own use | 134 | 53.4 | 117 | 46.6 |
|  | Individual: production for own use and sale | 27 | 69.2 | 12 | 30.8 |
|  | Individual: production for sale | 9 | 81.8 | 2 | 18.2 |
|  | In farmers' association | 1 | 100.0 | - | - |
| >1ha | Individual: production for own use | 5 | 50.0 | 5 | 50.0 |
|  | Individual: production for own use and sale | 33 | 75.0 | 11 | 25.0 |
|  | Individual: production for sale | 59 | 95.2 | 3 | 4.8 |
|  | In farmers' association | 43 | 95.6 | 2 | 4.4 |

The sociodemographic and farming habit factors that were examined did not show a strong correlation with pesticide use (and nor with each other); the highest significant correlation (p < 0.05, Spearman’s Rho = 0.35) was with what growers thought about the risks of pesticide use (**Supporting Information S5**). However, the GBM model suggested that the best predictors for pesticide use in agricultural areas were if the respondents had consulted with an expert or were themselves trained agricultural experts (relative influence: 27.04, 84.4% of those who do versus 42.0% of those who do not consult with experts, or are expert themselves, used pesticides) and the respondents’ gender (relative influence: 18.41, 50.5% of females versus 82.1% of males used pesticides) (model accuracy: 0.79, AUC = 0.86, Sensitivity: 0.89, Specificity: 0.60) (**Figure 2, Supporting Information S6**).

**Figure 2.**
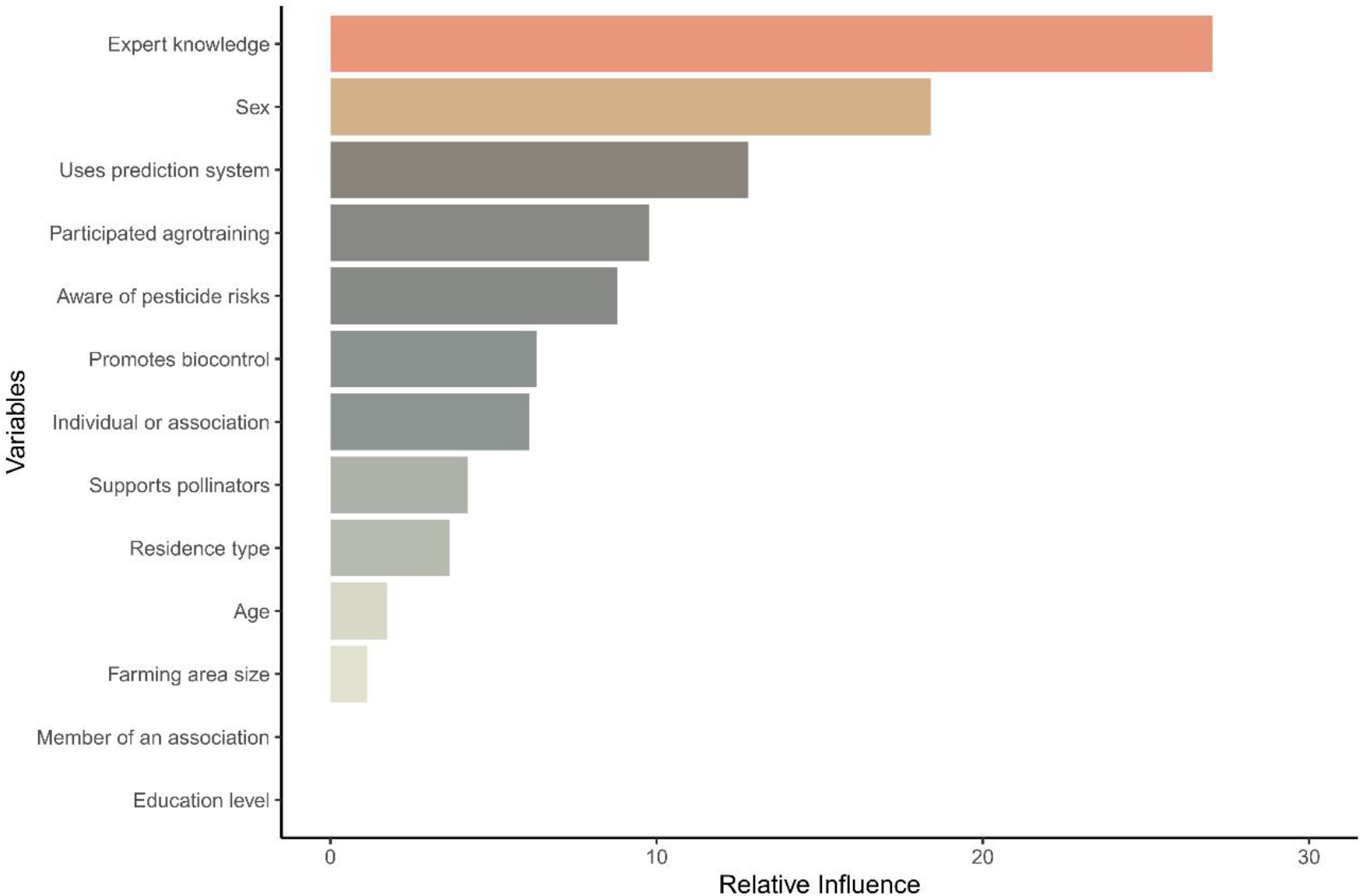
The relative influence of factors generated from the Gradient Boosting Machine (GBM) model for predicting pesticides used in farming areas.

Out of the 311 pesticide users, 243 (78.1%) felt that the use of insecticide was particularly indispensable for them. The main aspects that determined the choice of an insecticide were if they were harmless to humans, posed a low risk for bees and whether growers had previous experience with the product. The techniques by which the insecticide can be applied (such as if they can be used in flowering crops, if they can be used during daytime, or if they can be sprayed from the air) were the least important aspects to users (**Figure 3**).

**Figure 3.**
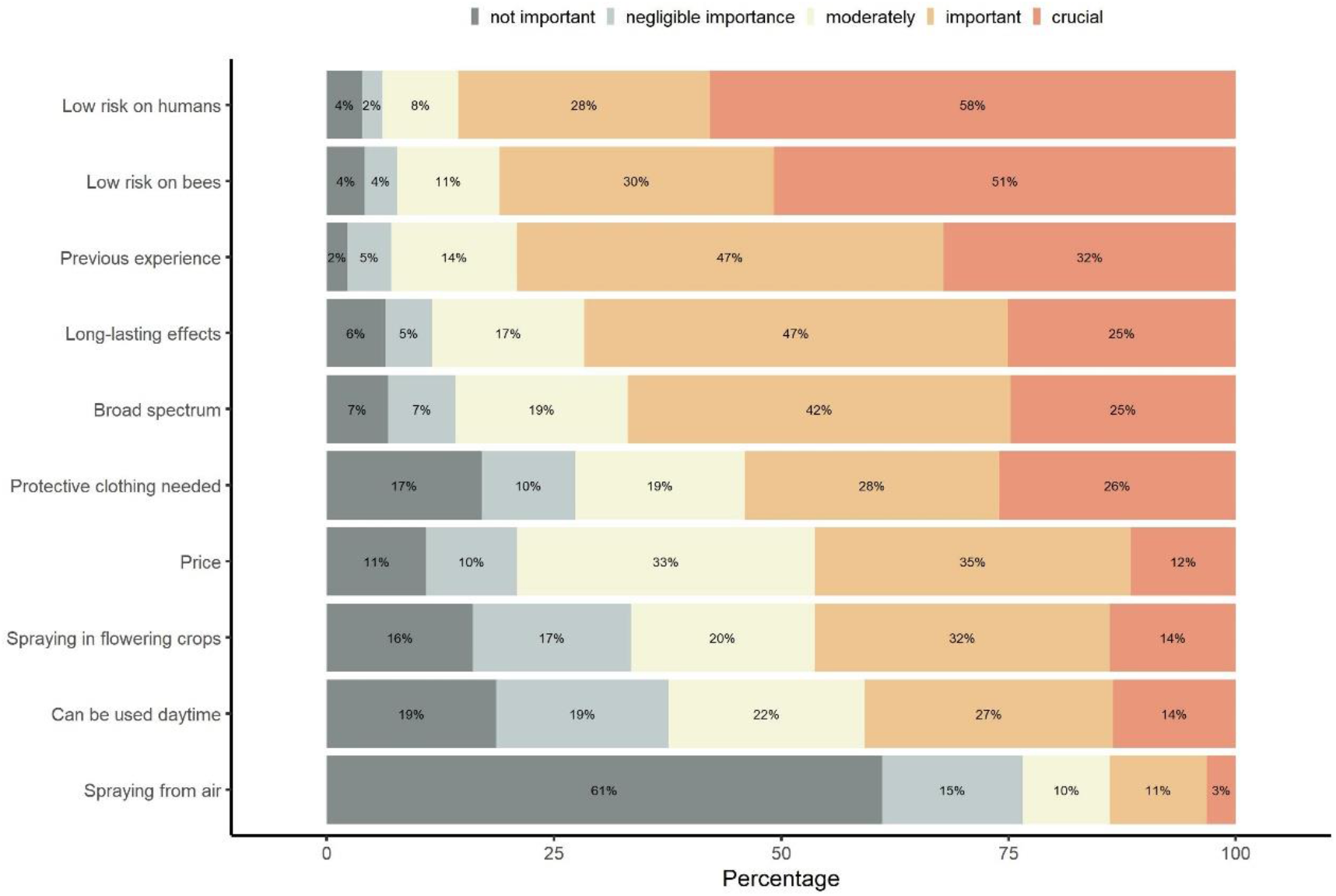
Relative frequency (%) of respondents’ opinions on how important different considerations are when choosing an insecticide. The response is colour-coded as follows: dark grey – not important, light grey – negligible importance, light yellow – moderately, light orange – important, dark orange – crucial.

Of those who used pesticides, 143 (46.0%) thought that banning neonicotinoids in the EU impacted their management practices, and 218 (70.1%) used at a minimum one acetamiprid-containing insecticides. Most users (55.2%) consider Mospilan (an acetamiprid-containing insecticide) indispensable. Of those who used Mospilan, 124 (56.11%) use it together with other pesticides, mostly with fungicides.

### 3.2 Plant protection habits and the protection of pollinators of garden owners

Of all questionnaire respondents, 302 (65.2%) had less than one hectare of land and 171 (56.6%) of home garden owners used pesticides on their land. The use of pesticides was considered acceptable by most garden owners, and 34.5% and 31.6% of them even thought it was important or crucially important, respectively. However, a significantly lower proportion of garden owners than of larger-scale farmers used pesticides (Chi-squared = 42.455, p-value <0.001) and a significantly higher proportion of home garden owners than of large-scale farmers believed that pesticide-free farming is achievable (Chi-squared = 3.593, p-value = 0.029).

The home garden owners who responded to our questionnaire specified that widespread use of pesticides, habitat loss due to agriculture and intensive agricultural production were the three most likely threats for wild pollinators (**Figure 4**), whilst they thought the appearance of invasive species was the least significant. Nonetheless, this factor was labelled as crucial by over half of the respondents (**Figure 4**). Of these garden owners, 259 (85.8%) recognise that widespread use of pesticides is a crucial problem for wild pollinators and 87.7% have heard that certain pesticides that are considered safe may also harm these insects. A significantly higher proportion of home garden owners than of large-scale farmers assumed that the conversion of agricultural production can slow down the depletion of pollinator populations (Chi-squared = 10.998, p-value = <0.001).

**Figure 4.**
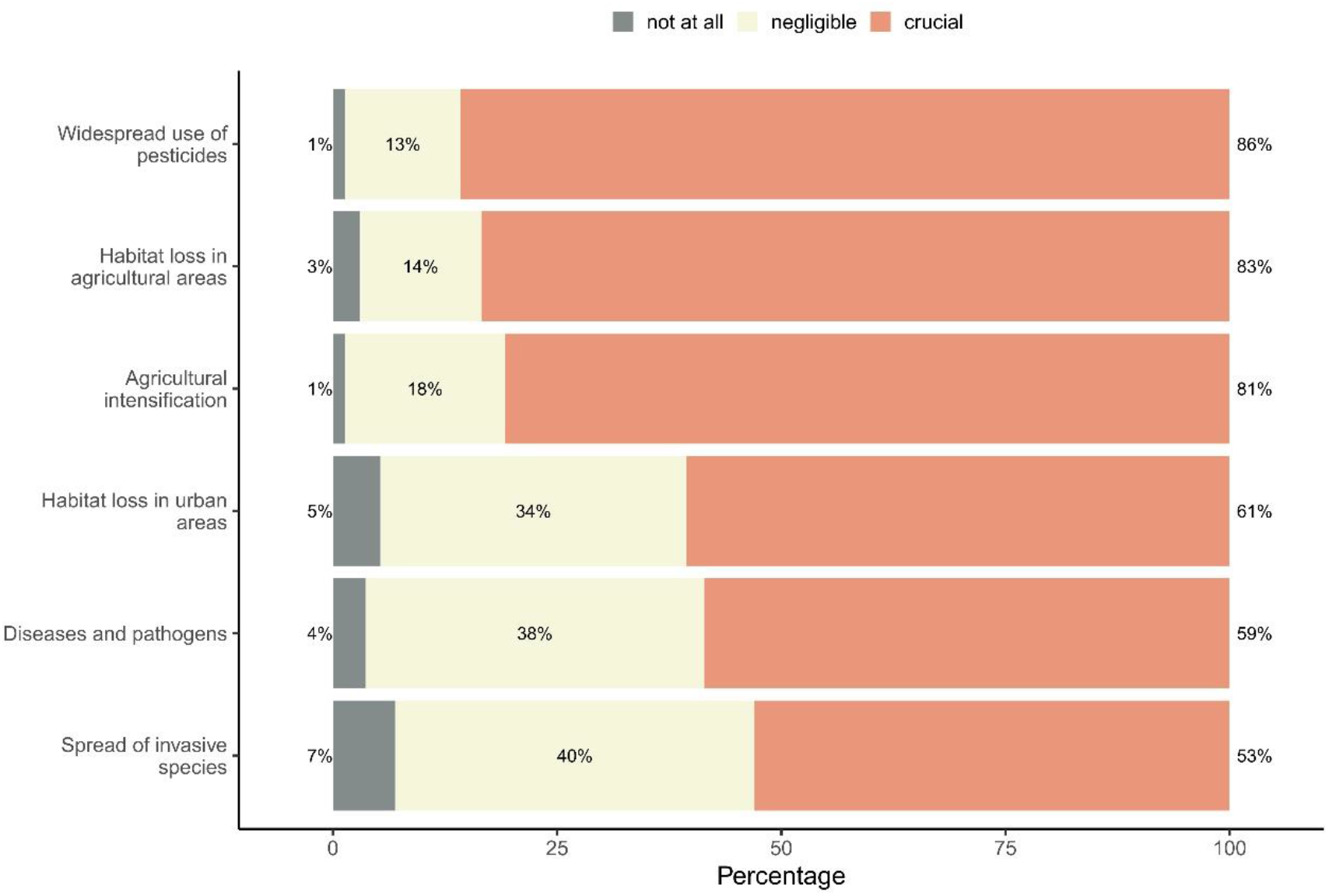
Relative frequency (%) of garden owners’ (with less than 1-hectare of land) opinions about the importance of factors that may threaten wild pollinators. The response is colour-coded as follows: grey – not at all, yellow – negligible, orange – crucial.

Of the garden owners, 81.1% carried out actions aimed at supporting wild pollinators. The examined sociodemographic and farming habits did not show a strong correlation with whether or not pollinators were supported (and neither did they with each other); yet the highest significant correlation with pollinator support was the growers’ pesticide use (p < 0.05, Spearman’s Rho = −0.22) (**Supporting Information S7**). The GBM model suggested that the best predictors for supporting pollinators were whether garden owners had promoted biocontrol (relative influence: 38.85) and garden owners’ age (relative influence: 26.34) (**Figure 5)**. This model had relatively high accuracy (0.82), and sensitivity (0.96) though only a moderate AUC (0.73), and very low specificity (0.18) (**Supporting Information S8**).

**Figure 5.**
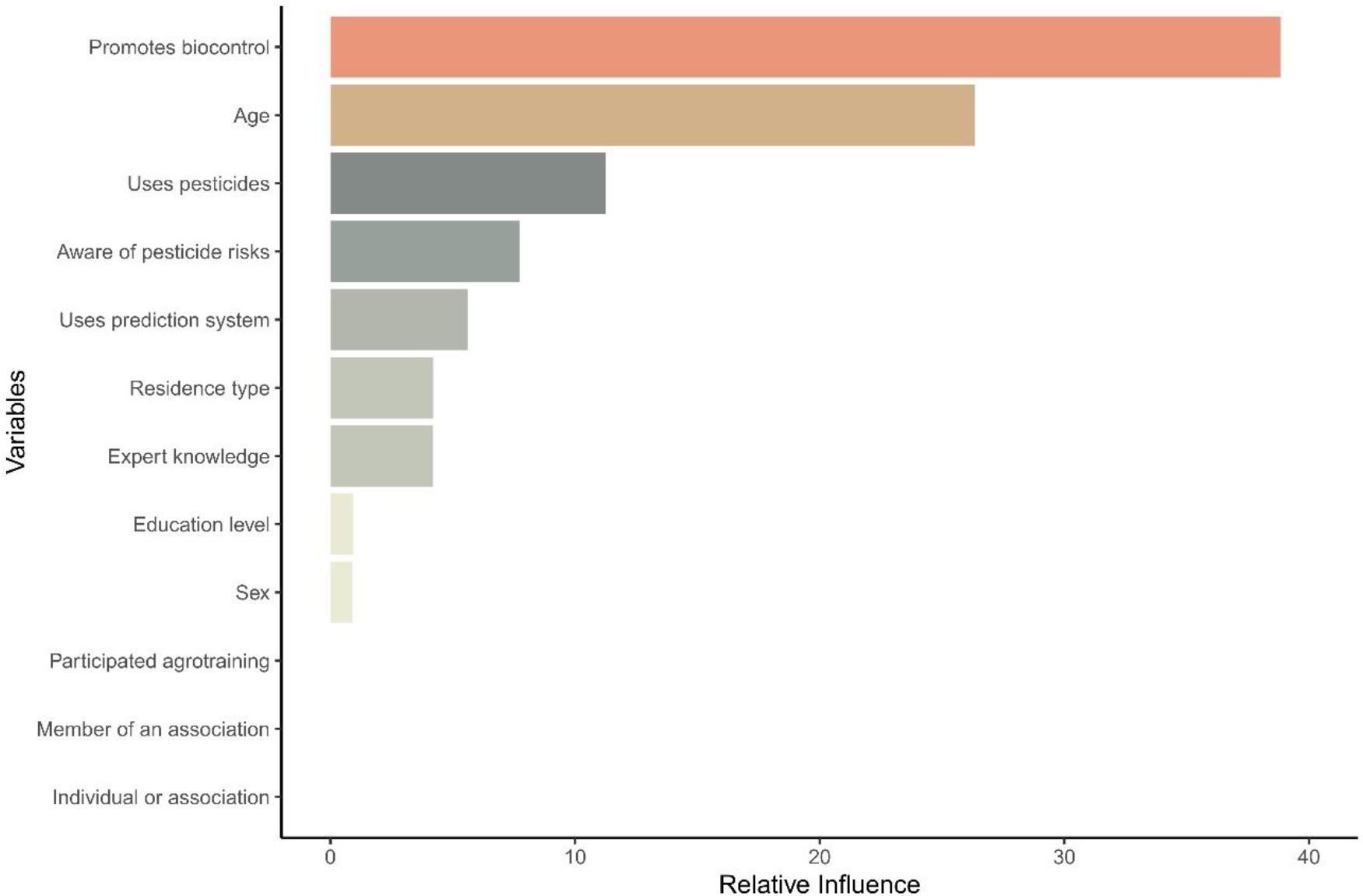
The relative influence of factors generated from the Gradient Boosting Machine (GBM) model for predicting whether or not garden owners support wild pollinators.

The proportion of those who supported pollinators was not significantly different between home garden owners and large-scale farmers (Chi-squared = 0.856, p-value = 0.178). A significantly higher proportion of pesticide-free garden owners supported pollinators than pesticide-using garden owners (Chi-squared = 13.159, p-value = <0.001).

Among home garden owners, the most common activities to support wild pollinators were to provide additional food sources (37.3%) (primarily by pollinator-friendly flowers) and natural habitat improvement (35.7%) (e.g. wildflower strips). Providing artificial habitat (19.9%) (e.g. bee hotels) and water (4.1%) were other forms of support. In some cases (3.1%), growers claimed they support pollinators without providing additional information. One respondent actively educated the neighbouring areas about the importance of wild pollinators and how to protect them.

## 4 DISCUSSION

In this study, we conducted an online survey in Hungary to investigate pesticide application practices of plant growers, particularly of home garden owners and their dependence on pesticides. Additionally, we also investigated the garden owners’ perspectives on environmental issues related to pesticides and their attitude to mitigating pollinator declines.

Supporting our first hypothesis, we found that almost half of those who completed our questionnaire claimed that general pesticide use is unavoidable in farming. This proportion was even higher amongst those who actively used pesticides.

We found that expert knowledge was the best predictor of whether pesticides were used in farming, and this was disproportionally important for large-scale farmers. Most of the respondents who usually consult an expert, or who are experts themselves with plant protection qualifications (e.g. plant doctor degree), use pesticides. Thus, farmers rely on (external) expert information for making decisions and embracing alternative pest management practices this expert advice may be essential for encouraging growers to move away from pesticide-based farming. The economic value of pollination ecosystem services ^59^ and the yield losses related to pollinator decline ^60^ may be the most important points to raise in addition to emphasising that maintaining pollinator populations requires drastic reduction or complete abandonment of pesticide use ^61,62^. However, farmers who grow crops that are not dependent on insect pollination and do not face the negative effects of their decline may be sceptical about the importance of this issue. Nonetheless, in our study, 40% of large-scale farmers personally observed pollinator declines and over 70% of them believed that transitional agriculture can mitigate pollinator declines. These results suggest that most growers are aware of the problem, yet their high level of dependency on pesticides implies a distrust or lack of knowledge of alternative methods. Also, for some crops no satisfactory management alternatives that protect yields are available. Environmental education, subsidising ecological management (e.g. agri-environment schemes supporting less intensive farming), and effective biodiversity offset schemes can play an important role, especially when combined with expert advice. However, the accessibility of this information varies from country to country ^63^, and so increasing the ease with which stakeholders can access this information is key in the transition process. Moreover, pressure from the agricultural chemical lobby and the distrust among agricultural advisers of alternative plant protection measures can strengthen market resistance ^64^, which can slow down the dissemination of ecologically friendly practices.

The second-best predictor of whether respondents used pesticides or not was gender. Despite genders being evenly distributed among respondents, almost twice as many men used pesticides as women. Indeed, in many respects, for instance, in eating habits (such as food-selecting behaviour), women are more health-conscious than men ^65^, which, we can speculate, may be reflected in differences in attitudes towards pesticide use (e.g. Wang et al. ^66^). Similar behavioural backgrounds may have created the emerging between-gender imbalance in our study.

Besides showing patterns of general pesticide use, our survey showed that the most important aspect for specifically choosing insecticides was the level of their effects on humans and bees. This suggests that most users were aware that insecticides can cause adverse, mostly sublethal, effects both in humans ^9^ and non-target insects ^67,68^. This was further underpinned by the large proportion of respondents (86.5%) who were aware that even insecticides labelled as harmless to insect pollinators can nevertheless have negative effects. Previous experience with a particular insecticide also influenced users’ choices. Repeatedly using well-known pesticides, however, may relax rigorous portioning habits which, in turn, may lead to insecticide overuse ^69^. This fixed choice may also lead to brand fidelity, which, consequently, may prevent experimenting with alternative, more environmentally friendly, pesticides.

Indeed, despite scientific advice calling for the banning of all neonicotinoids ^70^, this study showed that most respondents already experience the effects of the present ban on neonicotinoids and that they heavily depend on the use of acetamiprid, which is currently the only one freely available in the EU. This may lead to a higher demand for acetamiprid-containing insecticides among plant growers in the coming years ^71^. Acetamiprid, like all neonicotinoids, can persist in the tissues of treated plants ^45^ and its half-life can reach 450 days in soil ^72^ inducing sublethal effects in beneficial organisms ^71^, such as pollinators. On top of this, we also found that many of those plant growers who used acetamiprid-containing pesticides co-applied them in combination with other agrochemicals. Although concerns have been raised about the negative effects of cocktails of pesticides on the fitness of non-target insects (e.g. Gill et al. ^73^; Williamson and Wright ^74^), in our study the most extreme example was one home garden owner who used Mospilan along with seven additional fungicides. Based on our results, we can assume that a substantial proportion of Hungarian growers have not yet attempted to reduce insecticides. Similarly to when aiming to reduce pesticides at large, the publicising of relevant methodological advances or alternative technologies is likely to be critical to achieving a reduction in insecticides use and a transition to ecological-friendly farming.

### 4.1 Home gardens as ecological traps

Home gardens could be transformed into pesticide-free cultivation more quickly than larger-scale farming areas, but, according to our study, Hungarian garden owners seem to be reluctant of this conversion. Contrary to our expectations, pesticide use was widespread among gardeners, and almost all respondents who considered that the issue of pesticides causing harm to wild pollinators was unimportant themselves used pesticides. Even those garden owners who acknowledged that the widespread use of pesticides was a crucial problem for pollinators and have heard that certain pesticides considered safe may also be harmful to wild pollinators kept using them. Hence, our second hypothesis was not supported. Although a significantly greater proportion of home gardeners than of large-scale farmers believed that pesticide-free farming is achievable. Only 43.4% of the garden owners who completed our questionnaire grow plants pesticide-free, and more than half of the garden owners who produce fruits and vegetables for themselves and are not profit-oriented, use pesticides. These numbers are alarming and suggest that despite the known negative effects of pesticide use and the potential benefits of pesticide-free management, garden owners favour conventional approaches including the use of pesticides. The proportion of pesticide-free gardeners is similar to that found in Austria and Poland (pesticide-free: 41.0-51.7%) ^75^ among small-scale gardeners. In another survey conducted in the UK, only 30% of small gardeners did not use pesticides ^76^. However, the comparability of these results may be hampered by the differences in the definition of a ‘home garden’ among surveys.

The majority of those who completed the questionnaire supported pollinators. Our model indicated that whether or not one promoted biocontrol was the best predictor of whether a garden owner also supported wild pollinators. However, due to the skew in number towards pollinator-supporting garden owners, the model specificity was low, making this prediction unreliable. A significantly higher proportion of pesticide-free garden owners supported pollinators than of those who used pesticides, supporting our third hypothesis. The most common means to support wild pollinators were to provide pollinator-friendly flowers and many respondents provided bee hotels as a means of support. These two approaches are probably widespread because in recent years pollinators (particularly wild bees) have become an increasingly important part of environmental education programs in the European Union ^77^, including Hungary (e.g. the annual ‘Pollinators day’ event). Yet, Schmied et al. ^78^ demonstrated that urbanised areas, whilst being safe habitats in some cases (e.g. Theodorou et al. ^41^), can also act as ecological traps (e.g. Campioni et al. ^79^; Lehtonen et al. ^80^) for insects in other cases. Indeed, although home gardens lure insect pollinators, pesticides are used in many of those gardens, contaminating the nectar and pollen of flowers ^81^ which, in turn, can have deleterious effects on pollinators’ fitness. Thus, these non-pesticide-free gardens act as ecological traps for insect pollinators. For that reason, plant growers should be encouraged and motivated to produce their vegetables and fruits pesticide-free. Garden owners should be aware that to fully support pollinators in urban areas, pesticide use should be reduced or fully abandoned. Realising the potential benefits of urban gardens as biodiversity refuges, and the problems that pesticide use brings about for meeting this target, drive an increasing number of European countries to aim to ban plant protection products in private areas in addition to public areas ^7^. Home growers could also be discouraged from buying and using these products if they are removed from being freely available on supermarket shelves, as recently proposed in the UK ^82^.

### 4.2 Study limits and future perspectives

There are some limitations to our work. The respondents of the questionnaire were not chosen randomly. Our population is a subsample of those who were aware of the announcement and voluntarily took part in the study. The questionnaire could only be completed online, therefore, it had a lower chance to reach the eastern part of the country where there is a lower rate of internet access ^83^. Most of the large-scale agriculture takes place in Eastern Hungary, hence large-scale farmers may be underrepresented. Nonetheless, our questionnaire was completed by a sufficiently large number of people such that it should represent general plant protection habits and trends in pesticide use among Hungarian growers.

Our study could have provided further insights if more landscape and biodiversity variables had been available. However, asking for providing these may have been demanding for farmers and so reduced response rates. Thus, unfortunately, we had to compromise on this issue.

Furthermore, the generalisability of the result is limited by the fact that the survey was conducted only among Hungarian plant growers, although it does provide a useful case study in which results are not compounded by inter-country factors. Countries at a similar development level and using similar agricultural practices should also be involved in future studies to expand understanding of how pesticide use patterns and attitudes apply across a wider geographic scale.

### 4.3 Conclusions and future perspectives

Additional questions in similar studies could provide deeper insight into farmers’ practices, for instance, the chemical structure of pesticides used, whether they were synthetic or organic, or which organism were they used against.

One of the most pressing questions is whether home garden owners really want to convert to a completely pesticide-free plant growing. Focused research is needed to understand the willingness and motivations of garden owners for making this transition and why (if so) they prefer to continue conventional practices. Additionally, environmental education should establish the ecological foundations of pollinator-friendly gardens and promote their local and global benefits. In particular, demonstration gardens and demonstration farms should be set up which could demonstrate and teach pesticide-free farming to plant growers. Alternatively, incentives and direct subsidies could be provided to those who abandon the use of pesticides. Moreover, pesticide-free farming could be advocated through mobile applications and social recognition, or through granting ‘pollination friendly’ certification to home gardens. These gardens could also be involved in biomonitoring programs to further strengthen links between nature and garden owners.

However, at present, due to the unintentionally introduced pesticide pollution in gardens ^45^, not even the exact magnitude of exposure of pollinators to chemical pesticides can be assessed. Therefore, future research should focus on this invisible contamination and its effects on the assemblages of garden insects. From the side of decision-makers, to deal with this issue, clear labelling practices should be requested from suppliers to indicate whether or not products have come from pesticide-free farms (e.g. Ecolabel Index ^84^.). The amount of freely available plant protection products should be reduced, particularly those available in supermarkets, to discourage direct and indirect pesticide pollution in the gardens.

Driving large-scale farmers and home garden owners toward pesticide-free farming however may need different approaches. Whereas large scale farmers mostly rely on expert advice, and therefore advisers should inform them about pesticide-free practices, home gardeners may more heavily rely on conventional information channels (such as social media and personal networks).

The attitude of Hungarian plant growers can provide a general insight into the viewpoint of other Central and Eastern European countries and similar surveys would be needed across Europe. Since survey approaches, similar to ours, through directed questions about pesticide use habits, help us to better understand plant growers’ motivations, we hope this survey proves to be useful as an example for further online questionnaires. The information gained then can help to find solutions towards a pesticide-free future.

## ACKNOWLEDGEMENTS

We are grateful to Zoltán Tóth for pointing us towards conducting a questionnaire-based study. We also thank Marco Ferrante and Nick Littlewood for their comments on the manuscript. Zsófa Varga-Szilay was supported by the ÚNKP20-3 New National Excellence Program of the Ministry for Innovation and Technology from the source of the National Research, Development and Innovation Fund. Gábor Pozsgai was supported by the FCT-UIDP/00329/2020 funding and the research pluriannual funds for cE3c (FCT-UIDB/00329/2020).

## AUTHOR CONTRIBUTIONS

Zsófia Varga-Szilay conceived and designed the study. The questionnaire was created and data were collected by Zsófia Varga-Szilay. Analysis was performed by Zsófia Varga-Szilay and Gábor Pozsgai. The manuscript was written by Zsófia Varga-Szilay and Gábor Pozsgai. Both authors read and approved the final manuscript.

## DATA AND CODE AVAILABILITY STATEMENT

The data and the underlying computer code are available in the GitHub repository https://github.com/zsvargaszilay/pesticide_questionnaire

