## Supporting Information for "Plant growers’ environmental consciousness may not be enough to mitigate pollinator declines: a questionnaire-based case study in Hungary"

Pesticide use and pollinator support in Hungarian plant growers

Zsófia Varga-Szilay[1*], Gábor Pozsgai[2]

[2]cE3c – Centre for Ecology, Evolution and Environmental Changes/Azorean Biodiversity Group, CHANGE – Global Change and Sustainability Institute, Departamento de Ciências e Engenharia do Ambiente, Universidade dos Açores, Açores, Portugal,, https://orcid.org/0000-0002-2300-6558

**This file includes:**

Questions of the online survey S1

Tables S2, S3, S4

Figures S5, S6, S7, S8

**S1 Questions of the online survey (translated from Hungarian). Multiple choices were allowed (*). Textual response (†).**

1. Provide your gender.

   Male/Female/I don't want to answer.

2. Provide your age.

   8-25/26-35/36-45/46-55/56-65/over 65.

3. What is your highest level of education?

   Elementary/Middle/High/Postgraduate.

4. What is the type of your residence?

   Farmland/Countryside/Town/Major city/Capital.

5. In which Hungarian county do you farm (*)? Please enter the name of the settlement (†).

   19 Hungarian counties listed by name.

6. On what area do you farm?

   <1 ha /1-9.9 ha/10-29.9 ha/30-49.9 ha/50-99.9 ha/100-299.9 ha/300-499.9 ha/500-999.9 ha/≥ 1.000 ha.

7. Do you farm as a private individual or within a farmers' association?

   Private individual: production for own use/Private individual: production for own use and sale/Private individual: production for sale/In farmers' association.

8. Is your farm a conventional farm or an organic farm?

   Conventional farming/Organic farming.

9. Do you farm in an open field and/or in the greenhouse (*)?

   Field/Greenhouse.

10. What type(s) of cultivated plant do you grow (*)?

    Vegetables/Fruits/Grapes/Root vegetables/Tuberous plants/Cereals/Legumes/Oil seeds/Fodder/Spices, herbs, ornamental, other/Other industrial crops.

11. Do you consult with a plant doctor?

    Yes/No/No, because I am a plant doctor.

12. Do you consult with a plant protection expert?

    Yes, always/Yes, rarely/No/No, because I have the qualification.

13. Are you a member of a farming association?

    Yes/No.

14. Have you ever participated in an agrotraining of Integrated Pest Management (IPM)?

    Yes/No.

15. Do you use any forecasting system (e.g. pheromone traps) to track pest populations?

    Yes/No.

16. Do you support beneficial insects (e.g. natural enemies) to promote the biocontrol of weeds, pests and pathogens?

    Yes/No.

17. Do you use pesticide(s)?

    Yes/No.

18. What type of pesticide(s) do you use based on its/their application categorisation (*)?

    I. category/II. category/III. category/Organic products.

19. Have you reduced your pesticide use since 2014 when Integrated Pest Management was introduced?

    Yes/No.

20. How important do you think pesticide use is in farming?

    Indispensable/Important/Moderately important/Not important.

21. Do you keep a spray diary?

    Yes/No.

22. Do you keep a register book about pesticides used in our farming?

    Yes/No.

23. Do you use additive(s) with pesticides?

    Yes, always/Yes, not always/No.

24. Do you spray daytime in flowering cultures?

    Yes/No.

25. Can you imagine pesticide-free farming?

    Yes/No.

26. Do you feel that using insecticide(s) is indispensable for you?

    Yes/No.

27. How important the favourable price is for you when you choose an insecticide?

    Crucial/Important/Moderately/Negligible importance/Not important.

28. How important the broad spectrum effect is for you when you choose an insecticide?

    Crucial/Important/Moderately/Negligible importance/Not important.

29. How important the long-lasting effect is for you when you choose an insecticide?

    Crucial/Important/Moderately/Negligible importance/Not important.

30. How important the previous experience with a particular product is for you when you choose an insecticide?

    Crucial/Important/Moderately/Negligible importance/Not important.

31. When choosing an insecticide, how important is it for you that it can be used daytime?

    Crucial/Important/Moderately/Negligible importance/Not important.

32. When choosing an insecticide, how important is it for you that it can be used in flowering crop?

    Crucial/Important/Moderately/Negligible importance/Not important.

33. When choosing an insecticide, how important is it for you that it can be sprayed from the air?

Crucial/Important/Moderately/Negligible importance/Not important.

34. When choosing an insecticide, how important is it for you that it has a low risk of harming bees?

    Crucial/Important/Moderately/Negligible importance/Not important.

35. When choosing an insecticide, how important is it for you that it has a low risk of harming humans?

    Crucial/Important/Moderately/Negligible importance/Not important.

36. When choosing an insecticide, how important is it for you that it can be used without protective clothing?

    Crucial/Important/Moderately/Negligible importance/Not important.

37. Would you (regularly) submit requests for emergency exemption for the insecticides you use?

    Yes, every year/Yes, not every year/No.

38. Has it affected your management practices that some neonicotinoids (e.g. tiacloprid) had been banned in the European Union?

    Yes/No.

39. Do you use any of the following acetamiprid-containing insecticides (*, †)?

    Aceptorro/Apis/Artiler/Autentic/Careo/Gazelle/Inazuma/Los Ovados/Mospilan/Provado care/Rafting/Spilan/Other/Not.

40. Do/did you use the Mospilan 20 SG acetamiprid-containing insecticide?

    Yes, currently/Yes, in the past/No.

41. For how long have you been using Mospilan 20 SG (†)? If this year is the first year you use Mospilan, please write number 1.

    Please enter a number.

42. Do you feel that using Mospilan 20 SG is indispensable for you?

    Yes/No.

43. Do you use Mospilan 20 SG together with other pesticide(s) (*, †)?

    Yes/No/And please enter the name of the pesticide.

44. Is it important for you that Mospilan 20 SG can be used daytime in flowering crops?

    Yes/No.

45. Have you submitted requests for emergency exemption for Mospilan 20 SG before 2021?

    Yes/No.

46. Have you submitted requests for emergency exemption for Mospilan 20 SG this year (2021)?

    Yes/No.

47. For what type of cultures have you submitted emergency exemptions for Mospilan 20 SG this year (2021) (*, †)?

    Beet/Onion/Wheat/Barley/Oat/Corn/Sunflower/Poppy/(Water)melon/Walnut/Soy/Other

48. Insect populations are declining in the last decades. Have you observed a decreasing trend in your surroundings?

    Yes/No.

49. To what level do you think the loss of habitat in agricultural areas threatens wild pollinators?

Crucial/Negligible/Not at all.

50. To what level do you think the loss of habitat in urban areas threatens wild pollinators?

    Crucial/Negligible/Not at all.

51. To what level do you think the widespread use of pesticides threatens wild pollinators?

    Crucial/Negligible/Not at all.

52. To what level do you think the agricultural intensification threatens wild pollinators?

    Crucial/Negligible/Not at all.

53. To what level do you think the spread of invasive species threatens wild pollinators?

    Crucial/Negligible/Not at all.

54. To what level do you think diseases and pathogens threaten wild pollinators?

    Crucial/Negligible/Not at all.

55. Do you think that the transformation of agricultural production (e.g. by Integrated Pest Management) can slow down the decline of pollinator populations?

    Yes/No.

56. Have you heard about that pesticides labelled as environmentally harmless can still negatively affect insect pollinators (like bumblebees)?

    Yes/No.

57. Have you observed unusual behaviour of bees in your farming areas or the surroundings (*)?

    Yes, on honeybee(s)/Yes, on bumblebee(s)/Yes, on other wild bee(s)/I have not paid attention/I have not noticed.

58. Do you find bumblebee nests in your farming area?

    Yes, regularly/Yes, this has happened before/I have not paid attention/No.

59. Do you use pollinators for your agricultural activity (*)?

    Yes, honeybees/Yes, wild bees/Yes, bumblebees/No.

60. Do you support wild pollinators in your farming areas or the surroundings?

    Yes/No.

61. How do you support wild pollinators (*, †)?

    Bee hotels/Wildflower strips/Pollinator-friendly flowers/Other.

**S2 Distribution of the types of cultivated plants among the study population. The third column shows the growing percentage of each type of plant cultivated by all respondents (n = 463). Note, that percentages sum up to more than 100% because respondents were allowed to indicate more than one type of cultivated plant in the questionnaire**

| Types of cultivated plants | n | % |
|---|---|---|
| Vegetables | 319 | 68.9 |
| Fruits | 308 | 66.5 |
| Grape | 174 | 37.6 |
| Root/tuberous plants | 173 | 37.4 |
| Cereals | 110 | 23.8 |
| Dried legumes | 102 | 22.0 |
| Oil seeds | 94 | 20.3 |
| Fodder | 76 | 16.4 |
| Spices, herbs, ornamental, other | 30 | 6.5 |
| Other industrial crops | 19 | 4.1 |

**S3 Farming habits of plant growers (n = 463)**

| | Yes | | No | |
|---|---|---|---|---|
| | n | % | n | % |
| Consulted with an expert | 275 | 59.4 | 188 | 40.6 |
| Member of farmers' association | 81 | 17.5 | 382 | 82.5 |
| Participated agrotraining | 186 | 40.2 | 227 | 59.8 |
| Uses prediction system | 181 | 39.1 | 282 | 60.9 |
| Promotes biocontrol | 370 | 79.9 | 93 | 20.1 |

**S4 Pesticide use habits of plant growers (n = 311)**

| | Yes | | No | |
|---|---|---|---|---|
| | n | % | n | % |
| Keeps spray diary | 189 | 60.8 | 122 | 39.2 |
| Keeps register book | 168 | 54.0 | 143 | 46.0 |
| Uses additives | 212 | 68.2 | 99 | 31.8 |
| Spraying daytime in flowering cultures | 67 | 21.5 | 244 | 78.5 |
| Submit requests for emergency exemption | 42 | 13.5 | 269 | 86.5 |

**S5 Correlation matrix of sociodemographic factors and plant protection habits of Hungarian plant growers (n = 463). Values showing Holm-adjusted p-values and colour depths indicate correlation strength**

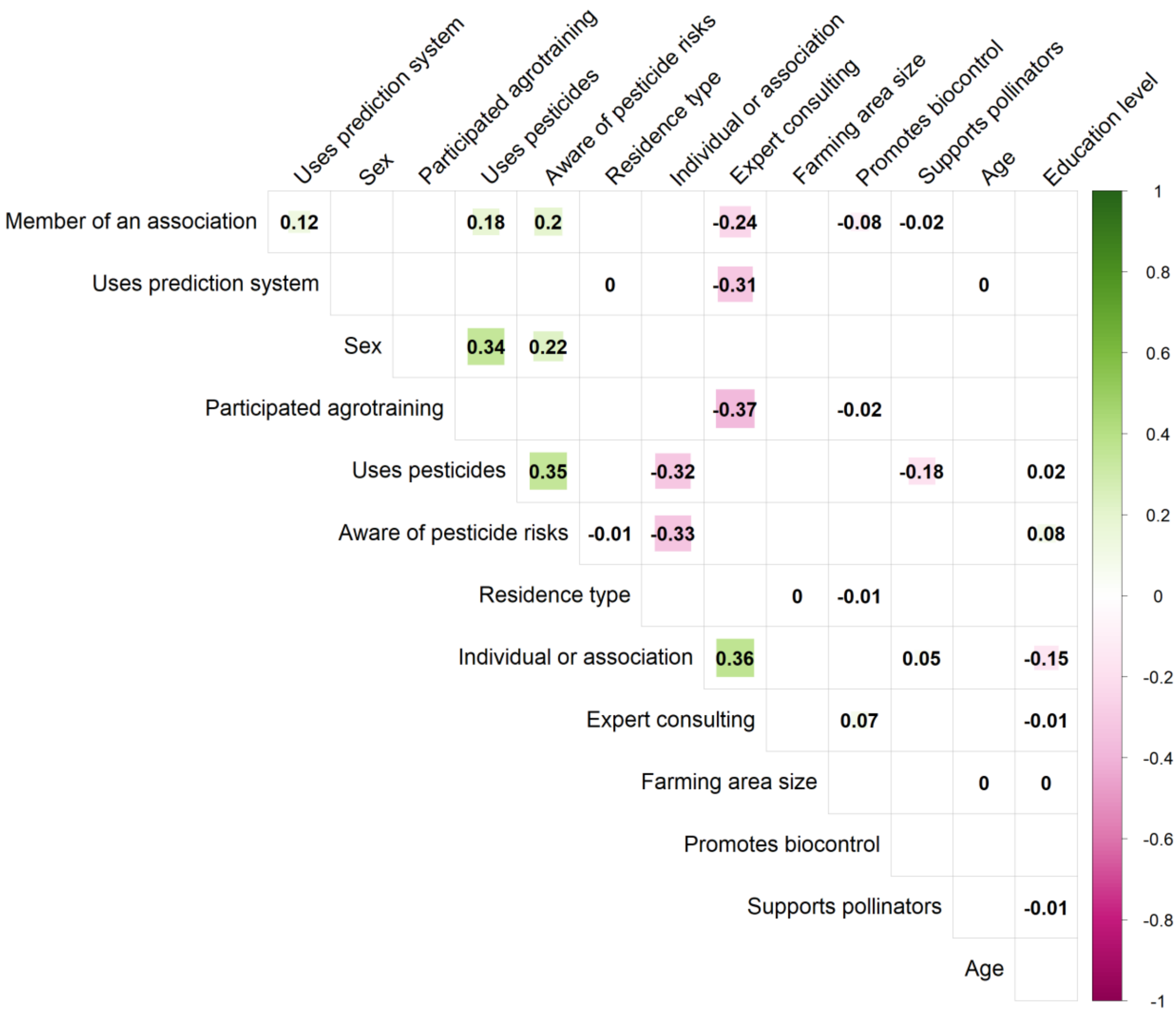

**S6 The performance evaluation of the Gradient Boosting Machine (GBM) model investigating the best predictors for pesticide use. Tuning parameter (top), ROC curve (bottom left), and Gain curve on test data (bottom right)**

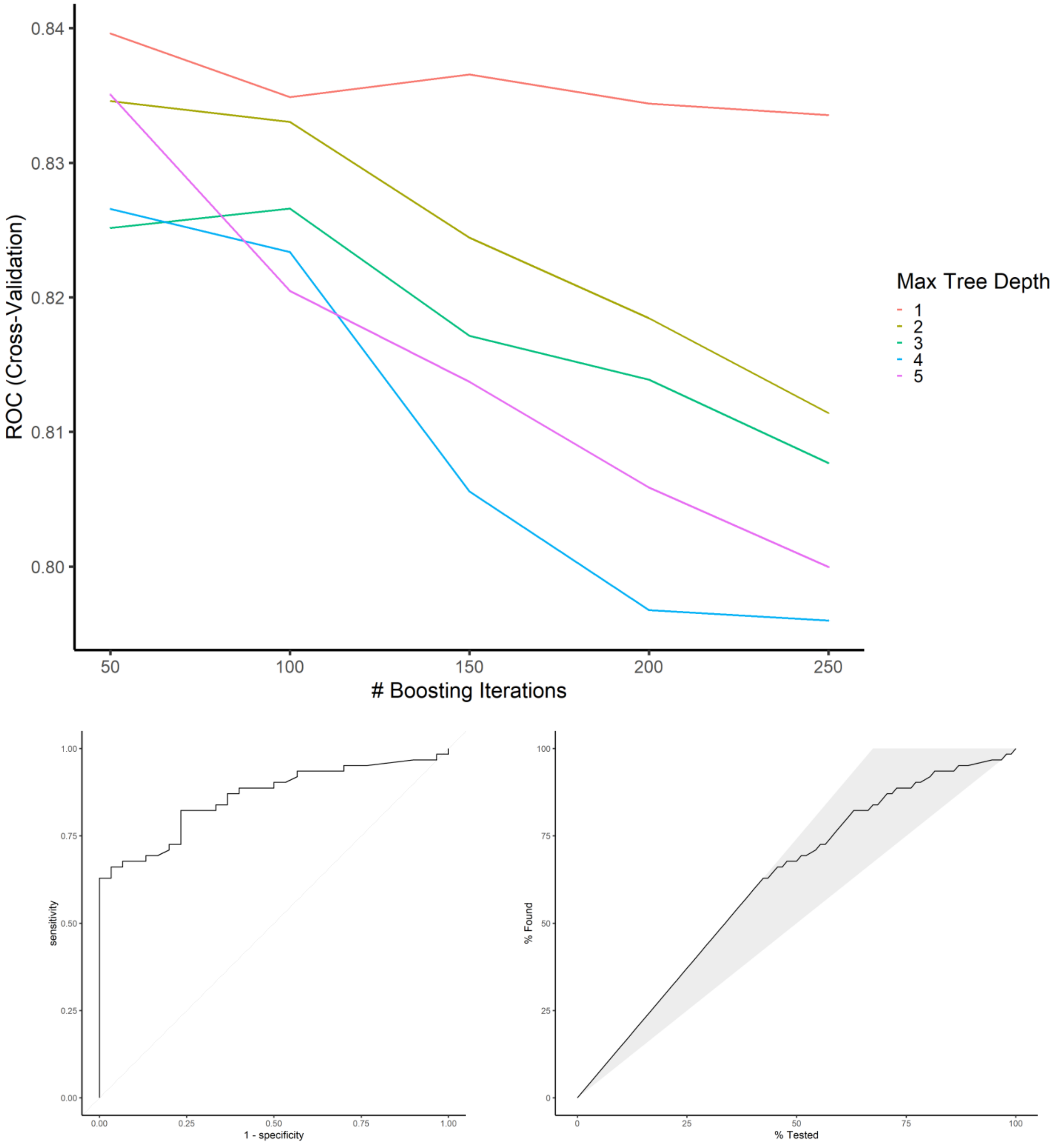

**S7 Correlation matrix of sociodemographic factors and plant protection habits of home garden owners (n = 302). Values showing Holm-adjusted p-values and colour depths indicate correlation strength**

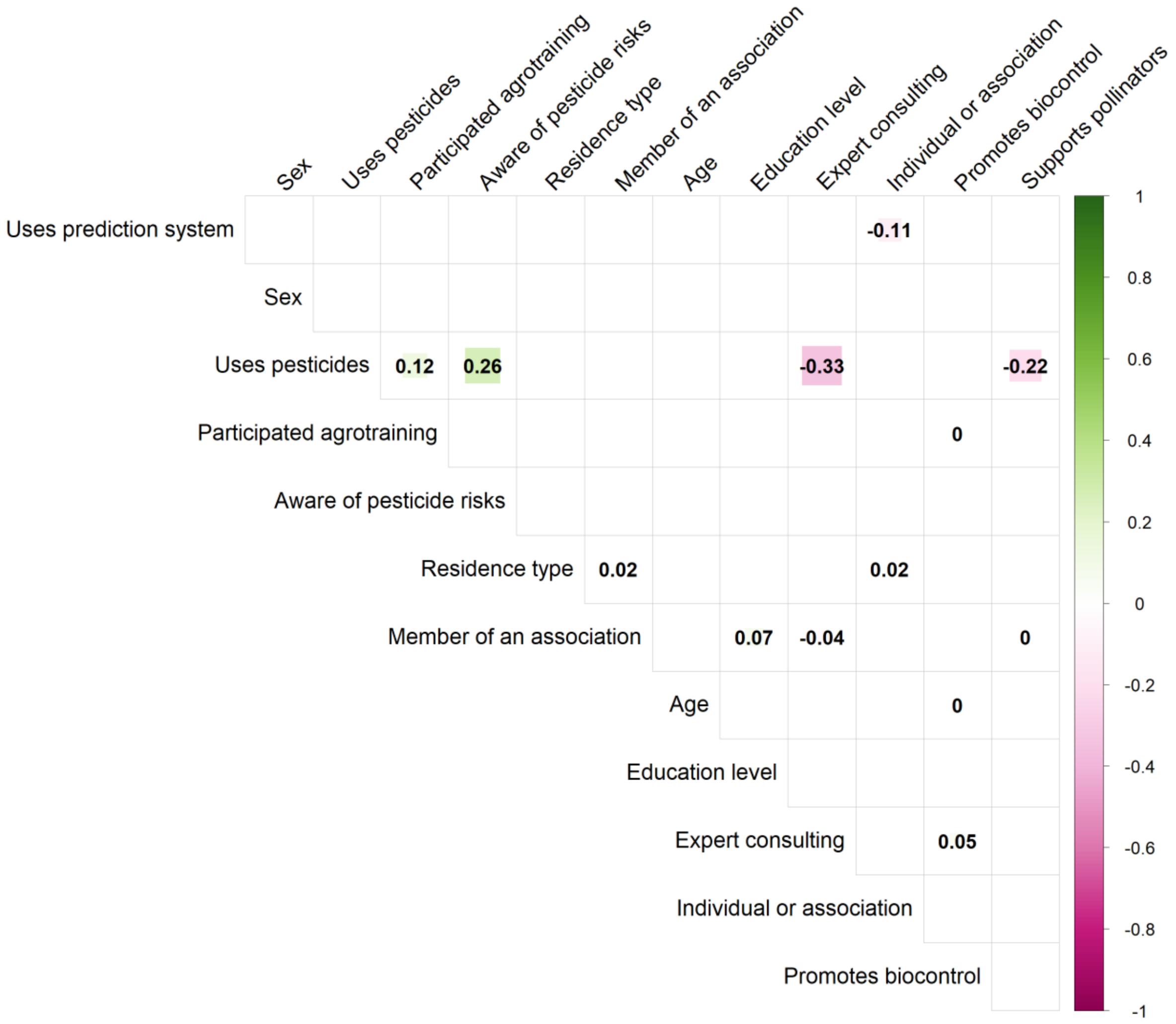

**S8 The performance evaluation of the Gradient Boosting Machine (GBM) model investigating the best predictors for supporting pollinators. Tuning parameter (top), ROC curve (bottom left), and Gain curve on test data (bottom righ)**

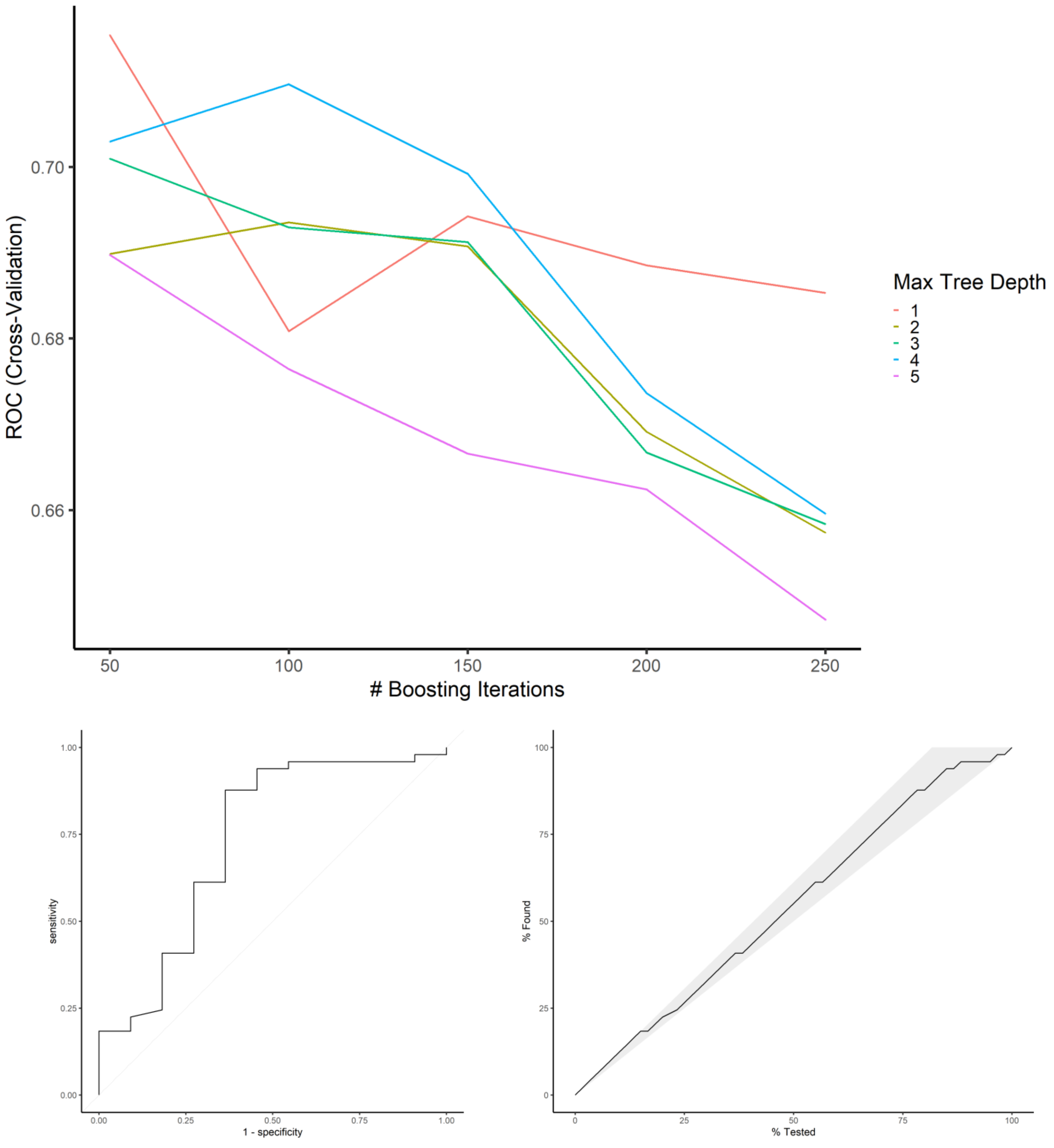
